# Elevated CD153 Expression on Aged T Follicular Helper Cells is Vital for B cell Responses

**DOI:** 10.1101/2023.03.17.533214

**Authors:** Alyssa L. Thomas, Joseph A. Wayman, Maha Almanan, Anthony T. Bejjani, Emily R. Miraldi, Claire A. Chougnet, David A. Hildeman

**Affiliations:** Department of Pediatrics, University of Cincinnati College of Medicine, Cincinnati, OH, United States; Division of Immunobiology of Cincinnati Children’s Hospital Medical Center, Cincinnati, OH, United States; Immunology Graduate Program, Cincinnati Children’s Hospital Medical Center and the University of Cincinnati College of Medicine, Cincinnati, OH, United States; Division of Biomedical Informatics, Cincinnati Children’s Hospital Medical Center, Cincinnati, OH, United States

## Abstract

Our recent data showed that an aberrant IL-10-producing T follicular helper population (Tfh10) accumulates dramatically with age and is associated with age-related declines in vaccine responsiveness. Through single cell gene expression and chromatin accessibility analysis of IL-10^+^ and IL-10^−^ memory CD4+ T cells from young and aged mice, we identified increased expression of CD153 on aged Tfh and Tfh10 cells. Mechanistically, we linked inflammaging (increased IL-6 levels) to elevated CD153 expression of Tfh cells through c-Maf. Surprisingly, blockade of CD153 in aged mice significantly reduced their vaccine-driven antibody response, which was associated with decreased expression of ICOS on antigen-specific Tfh cells. Combined, these data show that an IL-6/c-Maf/CD153 circuit is critical for maintaining ICOS expression. Thus, although overall Tfh-mediated B cell responses are reduced in the context of vaccines and aging, our data suggest that elevated expression of CD153 on Tfh cells potentiates the remaining Tfh function in aged mice.

## Introduction

Immune dysfunction in the elderly contributes to their diminished vaccine responses and increased prevalence and severity of infections (*1*). While aging negatively impacts nearly every compartment of the immune system, age-related deficiencies in CD4^+^ T cell function have been reported to substantially contribute to deficient immune responses in aging. Of importance, not only are there reduced numbers of naïve CD4^+^ T cells able to respond to new immunological challenges (*2*), but also age-acquired functional deficiencies such as decreased activation, aberrant cytokine production, decreased proliferative capacity, as well as increased frequency and number of immune-suppressive CD4^+^ T cell populations (*3-5*). Thus, there are multiple mechanisms that contribute to decreased immune responsiveness in the elderly and substantial efforts are being made to understand this mechanism with the hopes of reversing and/or restoring immune function in the elderly.

A subset of CD4^+^ T cells known as T follicular helper (Tfh) cells are responsible for providing aid to B cells and can promote the formation of germinal centers, immunoglobulin (Ig) class switching, and affinity maturation which are vital for producing adequate responses to both infections and vaccinations (*6*). Paradoxically, Tfh cells accumulate with age (*7*), while B cell responses decline with age, suggesting deficiencies in Tfh function in the aging immune system (*8-10*). One aspect of this reduced functionality centers on an inability to promote robust germinal centers in aged animals (*11, 12*). This results in defective maturation of germinal center (GC) Tfh cells and a reduced ability to help B cells (*13-15*). In addition to this defective maturation, our lab recently reported that a large fraction of Tfh cells that accumulate with age produce the anti-inflammatory cytokine IL-10, and the neutralization of IL-10R signaling restored B cell responses in aged mice (*7*).

Importantly, Tfh cells rely on many receptor ligand interactions to provide co-stimulatory signals to influence B cell responses. For example, CD154 (CD40L) and CD278 (inducible T-cell costimulatory, ICOS) both promote B cell survival and Ig class-switching (*16*). Interestingly, both CD40L and ICOS are also vital for Tfh differentiation and maintenance (*17-19*) and their expression is reduced on aged CD4^+^ T cells in mice and humans (*15, 20, 21*). In contrast, another co-stimulatory molecule, CD153 (CD30 ligand), has been reported to be elevated in senescent-associated memory CD44^+^ CD4^+^ T cells that also express the canonical Tfh markers PD1, CXCR5, and BCL6 (*22-24*). Further, both CD153 and CD30 are required for the accumulation of these senescent-associated cells and for autoantibody production in a mouse model of lupus nephritis (*25*). However, the role of CD153 in Tfh function is controversial as some studies suggest CD153 promotes B cell responses (*26, 27*), while others suggest that CD153 antagonizes them (*28, 29*).

Here, we investigated the role of CD153 in adaptive immune cell responses in aging. From our single-cell genomics analysis of the CD4^+^ memory T cell compartment (*30*), we identified *Tnfsf8* (encoding CD153) as a marker of Tfh whose expression increased with age. Notably, we verified that CD153 expression is increased on Tfh cells (PD1^+^ CXCR5^+^) with age. Further, our data suggests inflammaging, and in particular, elevated IL-6, is required to promote expression of CD153 on CD4^+^ T cells in aged mice.

Further, we provide evidence that IL-6 signaling through c-Maf promotes CD153 expression on CD4^+^ T cells. When assessed temporally after vaccination, expression of CD153 is transient on antigen-specific CD4^+^ T cells in young mice but sustained on the same cells in aged mice. Lastly, we show that CD153 neutralization during immunization leads to decreased antigen-specific, class-switched GC B cells and reduced antigen-specific serum antibody. Mechanistically, CD153 blockade reduces expression of ICOS on antigen-specific Tfh cells, which likely contributes to reduced antibody responses. In sum, although B cell responses in the elderly are defective, engagement of CD153 is critical for the remaining antibody producing capacity of B cells in the elderly.

## Materials and Methods

### Mice

Young (≤3 months) C57BL/6 mice were bred in house. Aged (≥18 months) mice were obtained from the National Institute of Aging (NIA) colony located at Charles River Laboratories (Wilmington, MA). Mice obtained from NIA were allowed to acclimate to our mouse facility for at least one week before experimentation. IL-6–deficient (IL-6 KO) mice on the C57BL/6 background were aged in-house. IL-10 GFP FoxP3 RFP mice were generated by crossing an IL-10–reporter (VertX) mice on the C57BL/6 background to a FoxP3-IRES-mRFP mice (The Jackson Laboratory) and were aged in house. All animal protocols were reviewed and approved by the Institutional Animal Care and Use Committee at the Cincinnati Children’s Hospital Research Foundation (IACUC 2019-0049).

### scRNAseq Data Acquisition and Analysis

Young (≤4 months) and old (≥18 months) IL-10 GFP x FoxP3 RFP mice were generated by crossing IL-10-reporter (Vertx) mice to FoxP3-IRES-mRFP mice and aged in house. All animal protocols were reviewed and approved by the Institutional Animal Care and Use Committee at the Cincinnati Children’s Hospital Research Foundation (IACUC 2019-0049). Spleens were harvested and crushed through 70-μm filters (BD Falcon) to generate single-cell suspensions. CD4^+^ memory T cells were enriched using the negative selection magnetic-activated cell sorting CD4^+^ T cell isolation kit II (Miltenyi Biotec, San Diego, CA). Enriched cells were stained with anti-CD4, anti-CD44, and anti-CD62L antibodies and sorted memory CD4^+^ T cells (CD4^+^ CD44^hi^ CD62L^lo^) that were either GFP^+^ (IL-10^+^) and GFP^-^ (IL-10^−^) by a FACSAria flow sorter (BD Biosciences).

scRNA-seq was analyzed as described in (*30*). In brief: Alignment, initial cell barcode filtering, and UMI counting was performed by Cell Ranger version 3.1.0 and their pre-built mm10 genome reference 2020-A. All downstream quality control and analyses was performed in R version 4.0.2, using workflows from Seurat v4.04 (*31*). Cell doublets were removed using DoubletFinder (*32*), setting the expected doublet rate to 5%. We retained quality cell barcodes with at least one UMI count from 500 genes, greater than 1,000 total UMI counts, and less than 15% of all UMIs corresponding to mitochondrial transcripts. We kept genes encoding proteins and lncRNAs on autosomal and X chromosomes detected in at least 20 cells. Pseudogenes as well as genes associated with ribosomal RNA contamination (*Gm42418* and *Ay036118*) and hemoglobin were removed. Initial, exploratory clustering of the combined samples revealed a population of primarily IL-10^−^ cells expressing Natural killer T (NKT) cell markers, which were removed from downstream analysis, given our focus on conventional CD4^+^ memory T cells. The final filtered UMI counts matrix contained 55,308 cells and 20,340 genes.

Expression was normalized in each cell by total UMI counts, scale to 10,000 total counts, and log-transformed (Seurat NormalizeData function). Within each biological replicate, we merged samples and selected a set of 2,000 genes with most variable expression across cells based on highest variance after a variance-stabilization transformation (“vst” option in Seurat FindVariableFeatures function). We mean-centered and variance-normalized expression of each highly variable gene across cells, regressing out variation due to total UMI counts (Seurat ScaleData function) and performed PCA dimension reduction, keeping the top 50 principal components (Seurat RunPCA function). Replicates were combined using Seurat’s anchor-based integration with the top 2,000 ranked variable genes across replicates (*33*). We scaled and performed PCA dimension reduction on the integrated dataset and identified clusters using the Louvain community detection algorithm (Seurat FindNeighbors and FindClusters functions). We computed the shared nearest neighbor (SNN) graph for 20 nearest neighbors using the top 50 principal components and identified clusters with resolution parameter 0.6. Initial clustering revealed a population of cells with high median total UMI counts and expressing markers of multiple T cell subsets. These cells were likely doublets and removed. Sample merging, replicate integration, and clustering were repeated as described above. The final T cell clusters were rigorously validated through cross-comparison to other relevant scRNA-seq studies as well as our (1) parallel single-cell chromatin accessibility (scATAC-seq) data and (2) bulk RNA-seq and ATAC-seq of flow-sorted Tfh populations (*30*). For final UMAP visualizations, we performed UMAP dimension reduction on the top 50 principal components (Seurat RunUMAP function). For heatmap visualization of pseudobulk gene expression patterns (Figure 1C), gene expression counts were summed per population and age group, normalized using DESeq2 VST transformation and z-scored. The raw and processed data are deposited in the GEO Database under accession GSE228668 (and a password for reviewers is cparwosghnuldcv.

**Figure 1:**
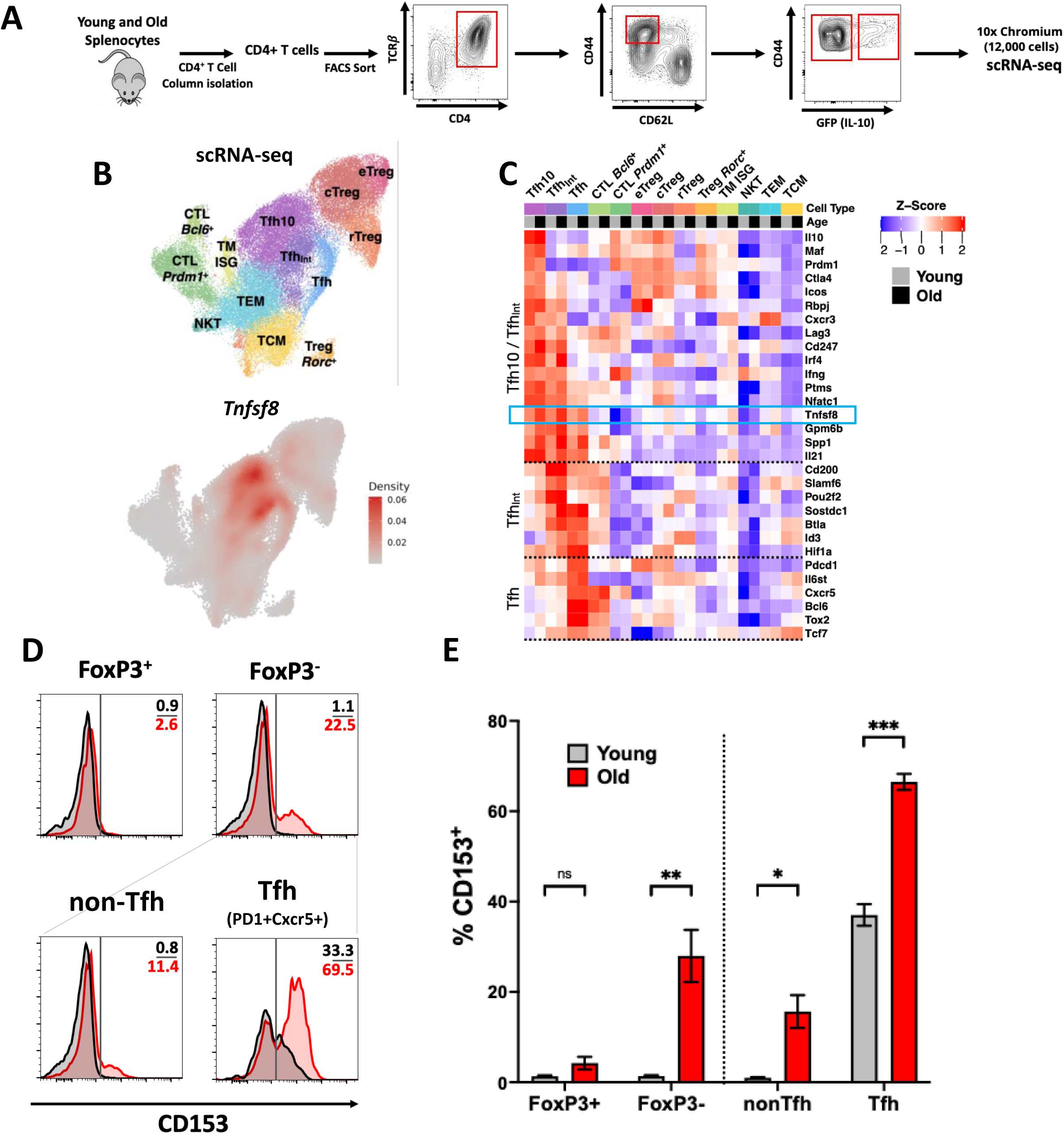
CD153 is a prominent marker of aged T follicular helper cells. **A)** IL-10^+^ and IL-10^−^CD4 memory T cells (TCRβ^+^ CD44^+^, CD62L^-^) young and aged IL-10 GFP x FoxP3 RFP mice were sorted and used for scRNAseq and nuclei were extracted for scATACseq. **B)** UMAP dimension reduction of scRNA-seq with subset annotation and density of *Tnfsf8* expression on the UMAP. **C)** Gene expression of selected markers across all cluster populations and age group. **D**,**E)** Splenocytes of young and aged C57BL/6 (n=4/group) were stimulated with P + I, stained with antibodies against TCRβ, CD8, FoxP3, IL-10, and CD153, and analyzed by flow cytometry. **D)** Representative histograms and **E)** graphs show the frequency of CD153^+^ cells in the CD4^+^ T cell compartment (TCRβ^+^ CD8^-^) (mean±SEM). \**P* ≤ 0.05, ** *P* ≤ 0.01, *** *P* ≤ 0.001, Student’s *t* test.

### Flow Cytometry

Single-cell spleen suspensions were generated by crushing spleens through 70-μm filters lysing red blood cells. A total of 3 × 10^6^ were plated per well, incubated with Fc block and surface-stained with a combination of the following antibodies: CD4, CD8α, TCRβ, CD44, PD1, CXCR5, CD30, CD153, ICOS, Fas, GL7, B220, CD19, and IgG1. To identify gp61-specific CD4^+^ T cells, splenocytes were stained with MHCII gp61 tetramer (National Institutes of Health). To identify NP-specific B cells, splenocytes were stained with APC-conjugated NP gifted by David Allman (University of Pennsylvania). For intracellular staining, cells were fixed and permeabilized using FoxP3/Transcription Factor Staining Buffer Set (eBioscience) and were then stained using antibodies for c-MAF and FoxP3. For cytokine staining splenocytes were first stimulated with PMA (25ng/ml) and ionomycin (0.5ug/ml) for 5 hours, with a brefeldin A and monensin block the final four hours. After 5 hours, cells were fixed, permeabilized, and intracellularly stained for IL-10 (Biolegend). Data were acquired on an LSR Fortessa flow cytometer (BD Biosciences) and analyzed using FlowJo software.

### Immunizations and Neutralization Experiments

For antigen-specific T cell tracking immunizations, mice were intraperitoneally immunized with 250μg gp61 (GLKGPDIYKGVYQFKSVEFDC) peptide conjugated to OVA (GenScript) mixed with 50% (v/v) alum (Thermo Fisher Scientific) and euthanized 10 days later. For antigen specific B cell tracking immunizations, mice were intraperitoneally immunized with 100μg NP-KLH (Biosearch Technologies) mixed with 50% (v/v) alum (Thermo Fisher Scientific) and euthanized 20 days after immunization. For CD30L neutralization experiments, mice were injected intraperitoneally with 500 μg of CD30L neutralizing antibody (CLONE M15-N297A), a generous gift from Amgen, Thousand Oaks, CA) on the day of immunization, and 4, 7 days after immunization.

### ELISAs

To measure NP-specific antibody titers, 96-well plates were coated with NP30-BSA (bovine serum albumin) (Biosearch Technologies) overnight at 4°C. Following blocking with 5% BSA for 1 hour at room temperature, serum samples were loaded into the plates with three dilutions per sample (1:27,000, 1:81,000, and 1:243,000) and incubated overnight at 4°C. Samples were washed and then incubated with a horseradish peroxidase-conjugated antibody reactive to mouse IgG1 (clone: PA1-74421; ThermoFisher Scientific) for 1 hour at room temperature. Samples were then washed and developed using ThermoScientific’s 1-Step Ultra TMB-ELISA solution and stopped after color change with 2N H_2_SO_4_. The plates were read at 450nm with an ELISA reader.

### In Vitro *IL-6 Stimulation*

CD4^+^ T cells were isolated from young WT mice spleens using Miltenyi’s CD4^+^ T cell isolation kit II and plated at 2.5 × 10^5^ cells per well in a 96 well flat-bottom plate. Cells were stimulated with 1 μg/ml anti-CD3 (Biolegend) and 1μg/ml anti-CD28 (BD Bioscience) in the presence or absence of 20ng/ml of recombinant mouse IL-6 (Biolegend) and/or 200ng/ml of nivalenol (Cayman Chemical) at 37°C. After 24 hours, CD153 and c-Maf expression was assessed via flow cytometry.

### Statistical Analysis

GraphPad Prism was used for graphing and statistical analysis. Statistical significance was determined by statistical tests indicated in each figure legend. The p values of <0.05 were considered to be significant and are indicated with the following nomenclature: * *p* < 0.05, ** *p* < 0.01, *** *p* < 0.001, ns = nonsignificant.

## Results

### CD153 is a prominent marker of aged Tfh cells

Our prior work showed that IL-10-producing CD4^+^ T cells accumulate with age, express markers of Tfh cells, and require the Tfh promoting cytokines IL-6 and IL-21 for their accrual, leading us to refer to them as Tfh10 cells (*7*). To further characterize the age-associated differences in IL-10-producing and non-producing T cells, we performed single-cell RNA sequencing (scRNAseq) of memory IL-10^+^ and IL-10^−^ CD4^+^ T cells from young and aged FoxP3 RFP x IL-10 GFP reporter mice (Figure 1A). We identified 13 clusters of CD4^+^ T memory cells (Figure 1B), including three populations of Tfh cells (*30*). Canonical Tfh cells (Tfh) were classified by expression of *Bcl6, Il21, Pdcd1*, and *Cxcr5*. Tfh10 cells expressed lower levels of *Bcl6*, but high levels of *Il21, Pdcd1, Cxcr5, Il10* and many other canonical Tfh signature genes. A third Tfh population expressed many genes in common with both Tfh and Tfh10 cells, with expression levels in between Tfh and Tfh10, and we designated them intermediate Tfh (Tfh_int_) (Figure 1C). Interestingly, there was a substantial increase in the expression of *Tnfsf8*, the gene that encodes CD153, within the three Tfh populations compared to all other memory CD4^+^ T cell populations, especially within aged cells (Figure 1C). To confirm these findings, we examined CD153 expression by flow cytometry across multiple CD4^+^ T cell populations in young and aged mice (Suppl. Figure 1A, B). Consistent with the scRNAseq data, CD153 was minimally expressed on FoxP3^+^ Tregs, while the majority of FoxP3^-^ CD4^+^ T cells that expressed CD153 were Tfh cells (Figure 1D). Further, the frequency of CD153-expressing cells increased substantially with age (Figure 1E). Similar differences existed when CD153 expression was compared in young versus old memory subsets (Suppl. Figure 2). Thus, these data show that CD153 expression is more prevalent in the Tfh populations, particularly in aging.

### CD153 expression is particularly increased on IL-10-producing T cells

Our scRNAseq data showed that CD153 mRNA expression was highest on IL-10-producing Tfh cells (Tfh10 cells). To determine whether this difference was observed at the protein level, young and aged mouse splenocytes were stimulated *ex vivo* with PMA and Ionomycin for 5 hours to induce IL-10 production and CD153 levels were assessed by flow cytometry. Within Tfh cells, a higher percentage of IL-10^+^ Tfh cells expressed CD153 relative to IL-10-Tfh cells (Figure 2A). Similarly, within both FoxP3^+^ and FoxP3^-^ non Tfh cells, a higher frequency of IL-10^+^ cells expressed CD153 relative to their IL-10^−^ counterparts (Figure 2A, B). These data show that CD153 expression increases on Tfh and non-Tfh cells with age, particularly on IL-10-expressing T cells.

**Figure 2:**
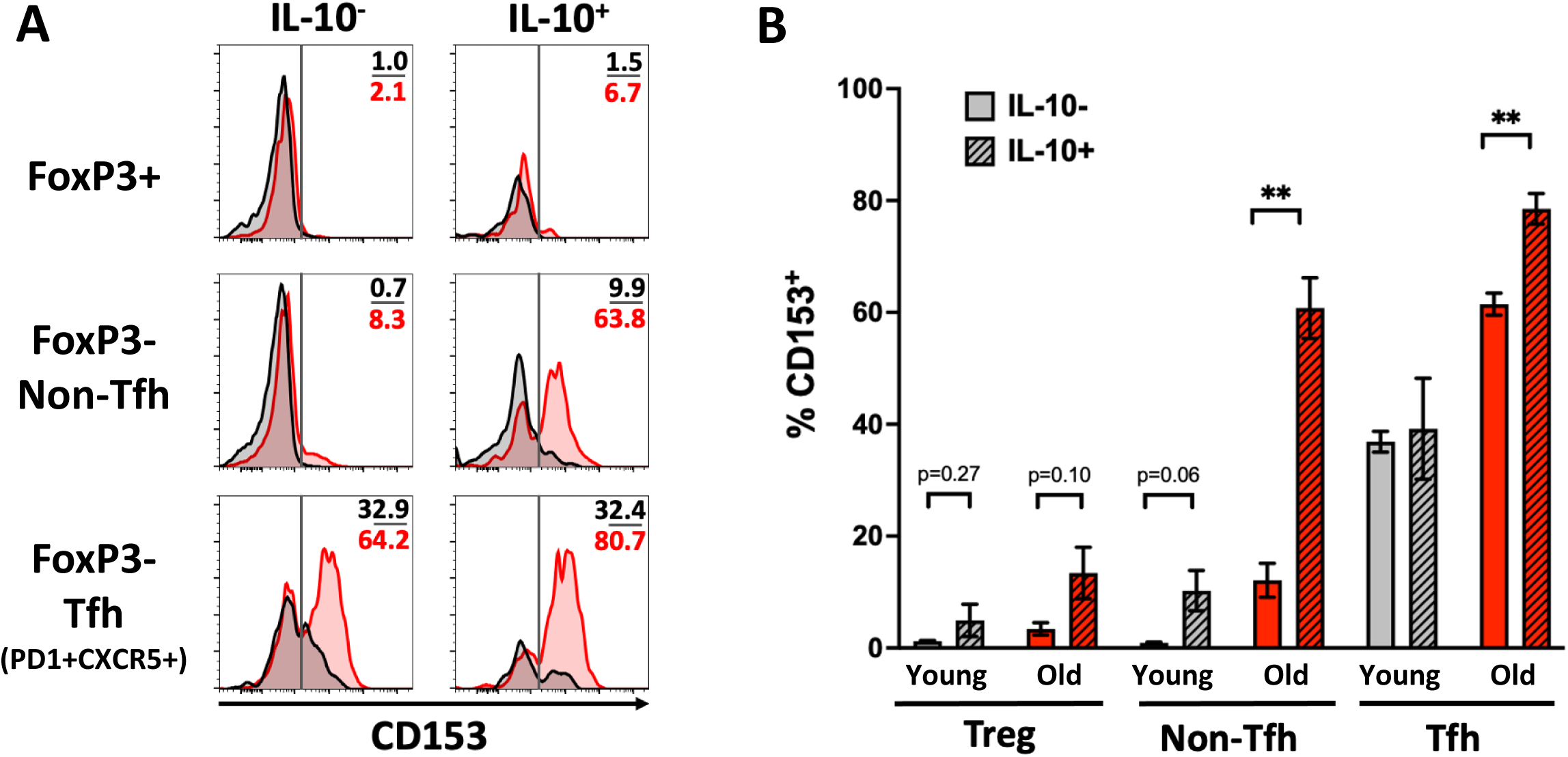
CD153 expression is particularly increased on IL-10-producing T cells. **A-B)** Splenocytes of young and aged C57BL/6 (n=4/group) were stimulated with P + I, stained with antibodies against TCRβ, CD8, FoxP3, IL-10, and CD153, and analyzed by flow cytometry. Representative histograms and graphs show CD153^+^ expression in the CD4 T cell compartment (TCRβ^+^ CD8^-^) based off their IL-10-producing status (mean±SEM). \**P* ≤ 0.05, ** *P* ≤ 0.01, *** *P* ≤ 0.001, Student’s *t* test.

### c-Maf, an important Tfh transcription factor, is elevated in aged Tfh and Tfh10 cell populations

We next aimed to understand mechanisms controlling Tfh function in aged mice. Several lines of evidence suggest that c-Maf is a key regulator of Tfh10 and might regulate *Tfnsf8* expression in Tfh and Tfh10 cells. Across the memory T cell populations, expression of the transcription factor *Maf* was highest in Tfh10 cells and increased with age in multiple populations, including the other Tfh/Tfh-like populations, as well as *Bcl6*^+^ CTL (Figure 3A). Furthermore, computational predictions from our CD4^+^ memory T cell gene regulatory network (GRN) (*30*) predicted that protein c-Maf activity was highest in Tfh10 and age-elevated in Tfh populations. c-Maf is a known regulator of Tfh cell development and promotes Tfh-derived production of IL-4 and IL-21 (*34-36*). c-Maf is predicted, by our GRN, to regulate *Il10* in Tfh10 and is known to promote IL-10 production in other CD4^+^ T cell populations (*37*). Analysis of our scATAC-seq data (*30*) suggests that c-Maf might also regulate *Tfnsf8*. In the *Tfnsf8* locus, we identified Maf motif occurrences in accessible chromatin regions in Tfh10; one such region coincided with a bona fide c-Maf binding site in another CD4+ T cell population (Th17) by ChIP-seq (*38*) (Figure 3B, gray box). This regulatory prediction is also supported by c-Maf perturbation data (knockout followed by RNA-seq) in Tr1 cells (*37*). Thus, we examined the protein levels of c-Maf in Tfh and Tfh10 cells using flow cytometry. Consistent with our scRNAseq and gene regulatory network analyses, c-Maf increased in aged Tfh (Figure 3C) and Tfh10 (Figure 3D) cells compared to their younger counterparts.

**Figure 3:**
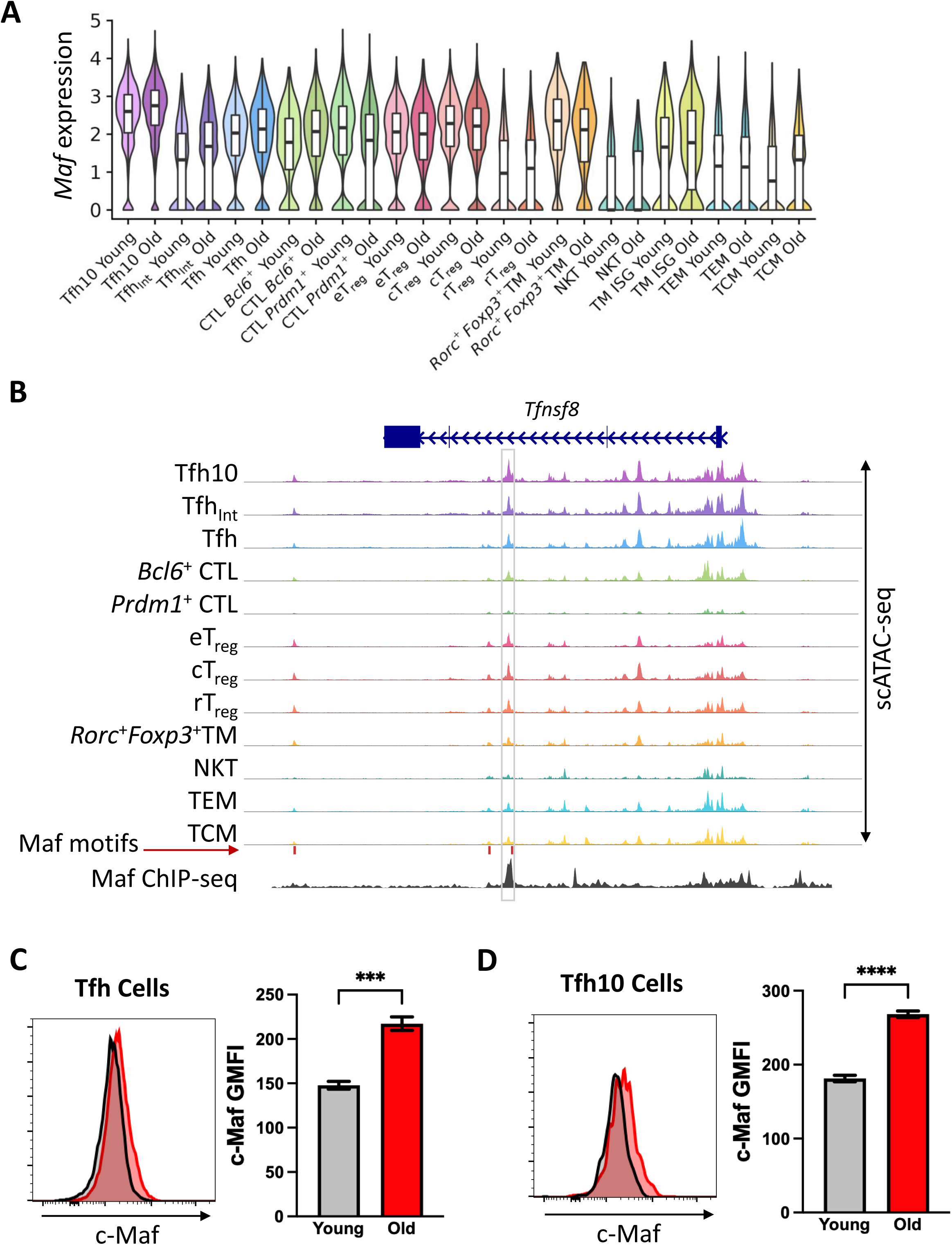
c-Maf, an important Tfh transcription factor, is elevated in aged Tfh and Tfh10 cell populations. **A)** *Maf* expression per cell in each CD4+ TM population (log-transformed transcripts per 10,000 transcripts). **B)** Epigenetic analysis the *Tfnsf8* locus (+/-10kb of the gene body): Pseudobulk scATAC-seq signal tracks are normalized to reads per million for each CD4+ TM population (young and old cells combined); Maf motif occurences (P<10^−5^, estimated by FIMO)(*56*); Maf binding (ChIP-seq signal) in Th17 cells. Flow cytometry analysis of cMaf expression in **C)** Tfh and **D)** Tfh10 cells in young (n=3) and aged (n=4) mice (mean±SEM). \**P* ≤ 0.05, ** *P* ≤ 0.01, *** *P* ≤ 0.001, Student’s *t* test.

### IL-6 controls CD153 expression through c-Maf

Given the collective evidence supporting a regulatory interaction between c-Maf and CD153, we next determined factors that promote expression of c-Maf and whether c-Maf regulates CD153 expression in Tfh and Tfh10. As we (*39*) and others (*40, 41*) have shown increased systemic IL-6 levels in aged mice, and IL-6 is known to induce c-Maf (*42*), we determined whether IL-6 controls c-Maf expression in Tfh and Tfh10 cells. In aged IL-6-deficient (IL6-KO) mice, the percent of Tfh and Tfh10 cells that express CD153 was reduced (Figure 4A). Further, IL-6 promoted c-Maf expression in both CD153^+^ and CD153^-^ Tfh and, to a lesser extent, Tfh10 cells (Figure 4B). Next, we determined whether c-Maf was required for IL-6 mediated upregulation of CD153. CD4^+^ T cells were isolated and stimulated *in vitro* for 24 hours with anti-CD3 and anti-CD28 in the presence or absence of recombinant mouse IL-6 with or without nivalenol, a small-molecule c-Maf inhibitor (*43*). As expected, IL-6 increased c-Maf (Suppl. Figure 3). Furthermore, IL-6 stimulation significantly increased the overall frequency of CD153^+^ cells and the per-cell level of CD153, in a c-Maf-dependent fashion (Figure 4B). Combined, these data provide a link between IL-6 mediated inflammaging and increased expression of CD153 via c-Maf.

**Figure 4:**
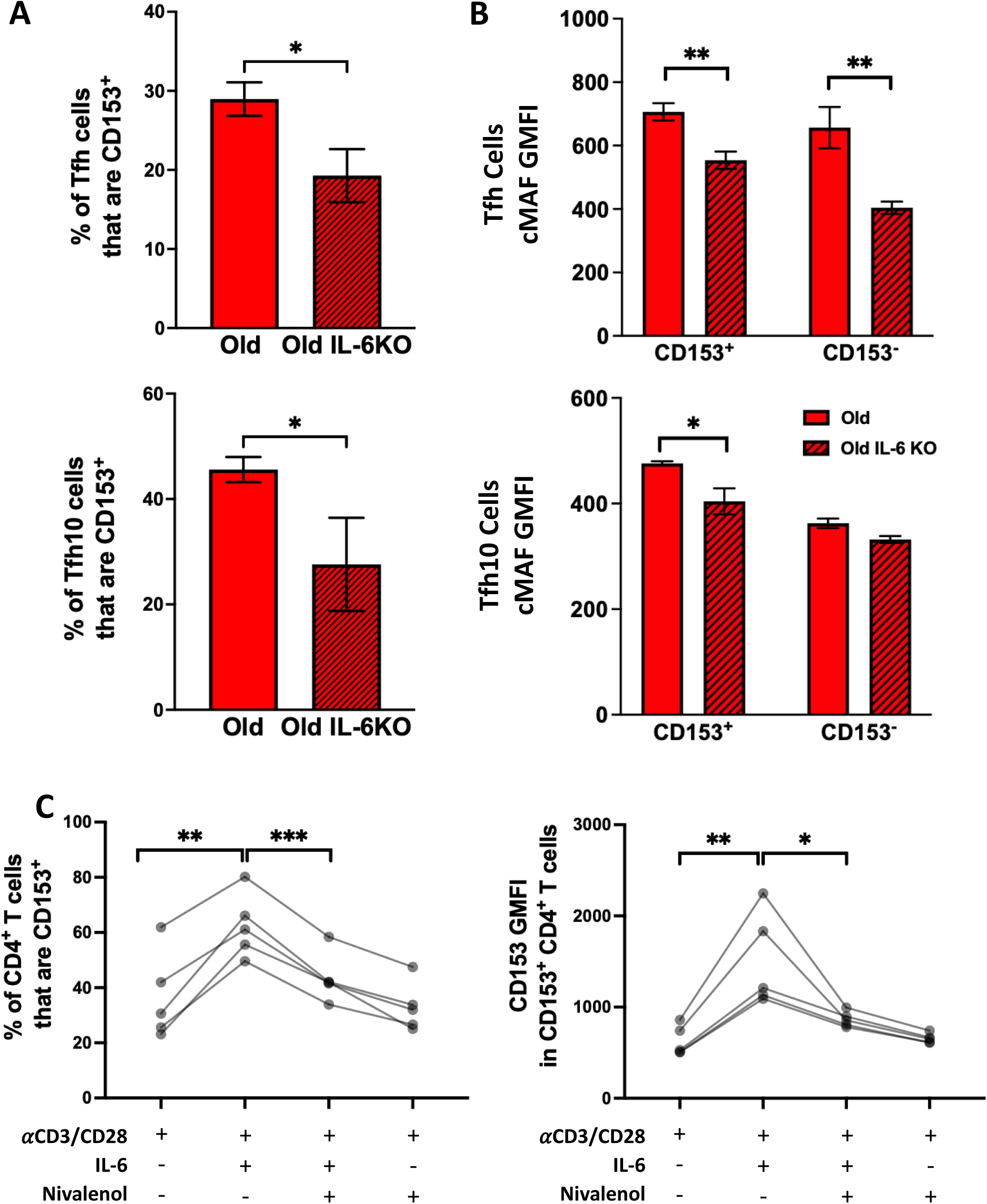
IL-6 controls CD153 expression through c-Maf. **A)** Aged matched WT and IL-6KO (n=7/group) mice were used to analyze the frequency of CD153^+^ Tfh and Tfh10 cells. **B)** Aged WT and IL-6KO (n=4/group) mice were used to measure c-Maf GMFI in Tfh and Tfh10 cells that were either CD153^+^ or CD153^-^. **c)** CD4^+^ T cells were isolated from young WT mice (n=5) and stimulated *in vitro* with *α*CD3/*α*CD28 in the presence or absence of IL-6 and nivalenol. GMFI of c-Maf **C)** Graph shows the frequency of CD153^+^ CD4^+^ T cells and gMFI of CD153 in those cells after 24 hours. Each line represents one mouse across treatment groups. \**P* ≤ 0.05, ** *P* ≤ 0.01, *** *P* ≤ 0.001, Student’s *t* test.

### CD153 expression is enhanced and persists on aged antigen specific Tfh cells after immunization

As the above experiments were performed on all Tfh and Tfh10 cells at steady state, we next examined whether CD153 expression was increased in antigen-specific Tfh and Tfh10 cells and whether it contributed to decreased vaccine responsiveness. We first developed a vaccination model that enabled us to track antigen-specific CD4^+^ T cells by conjugating LCMV immunodominant peptide gp61-80 to ovalbumin (gp61-OVA).

Groups of young and old C57Bl/6 mice were immunized with gp61-OVA and sacrificed 5, 10, and 20 days later. Gp61-specific (gp61-sp.) CD4+ T cells were assessed using flow cytometry (Figure 5A). Gp61-sp. CD4^+^ T cells emerged as early as 5 days post-immunization, peaked by day 10, and were decreased by day 20. Interestingly, aged mice had significantly more gp61-sp. CD4^+^ T cells than their younger counterparts on day 20 after immunization (Figure 5B). Further, within the gp61-sp. T cell population, there was a significant retention of Tfh (PD1^+^ CXCR5^+^) cells in aged mice which was evident on day 10 and persisted through at least day 20 after immunization (Figure 5C). A significant fraction of gp61-sp. Tfh cells expressed CD153 on days 5, 10, and 20 which was higher in aged relative to young mice (Figure 5D). Thus, CD153 is expressed on antigen-specific CD4+ T cells after immunization and is significantly increased with age.

**Figure 5:**
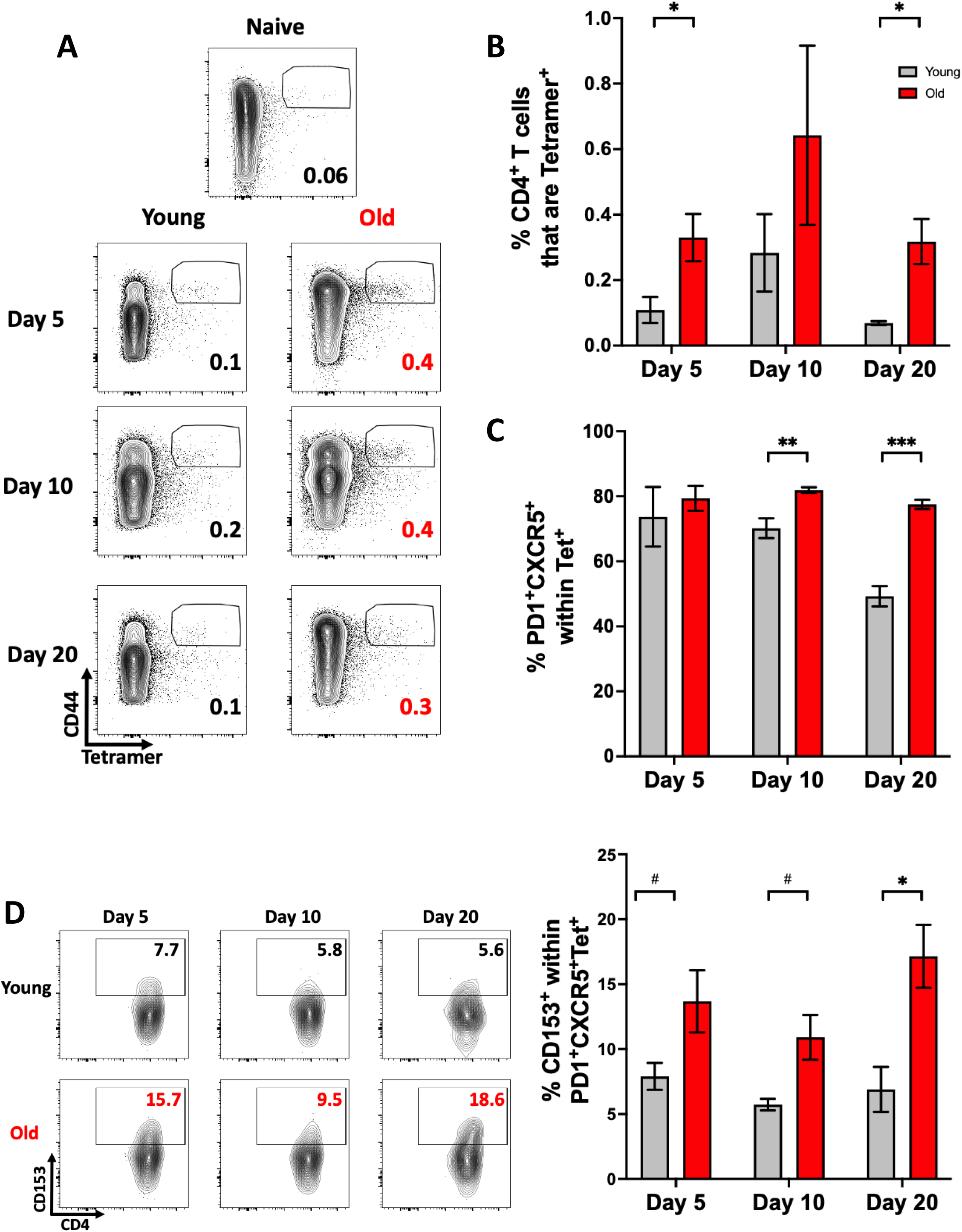
CD153 expression is enhanced and persists on aged Tfh cells after immunization. Young and aged WT (C57BL/6) mice were immunized with 250μg of gp61 conjugated to OVA (gp61-OVA) in alum and euthanized 5, 10, and 20 days later. (n=4/group) **A)** Plots show IA^b^ -gp61 tetramer staining in naïve and immunized young and old mice. Graphs show **B)** the frequency of gp61-specific total T cells and **C)** the frequency of Tfh cells (PD1^+^ CXCR5^+^) in the gp61-specifc T cell population (mean±SEM). **D)** Plots and graphs display the frequency of CD153^+^ gp61-specific Tfh cells (mean±SEM). # *P* ≤ 0.07, \**P* ≤ 0.05, ** *P* ≤ 0.01, *** *P* ≤ 0.001, Student’s *t* test.

### CD153 neutralization during immunization leads to decreased B cell responses

While Tfh cells are critical for productive B cell responses, the function of CD153 is controversial, some reports showing a stimulatory role (*26, 27*) while others suggest an inhibitory role (*28, 29*). We next tested the role of CD153 on vaccine-driven B cell responses. To track antigen-specific B cells, we utilized the classic nitrophenol-keyhole limpet hemocyanin (NP-KLH) (Figure 6A). Neutralization of CD153 with a non-depleting anti-CD153 antibody (a kind gift from Amgen) during NP-KLH immunization led to significantly decreased anti-NP IgG1 antibody levels in the serum on days 10 and 20 (Figure 6B). Further, flow cytometric analysis showed a decrease in GC B cells (Figure 6C), IgG1 class-switched GC B cells (Figure 6D), and significantly reduced frequency and number of NP-specific class switched (IgG1^+^) B cells (Figure 6E). These results show that CD153 promotes Tfh-dependent B cell responses in aged mice.

**Figure 6:**
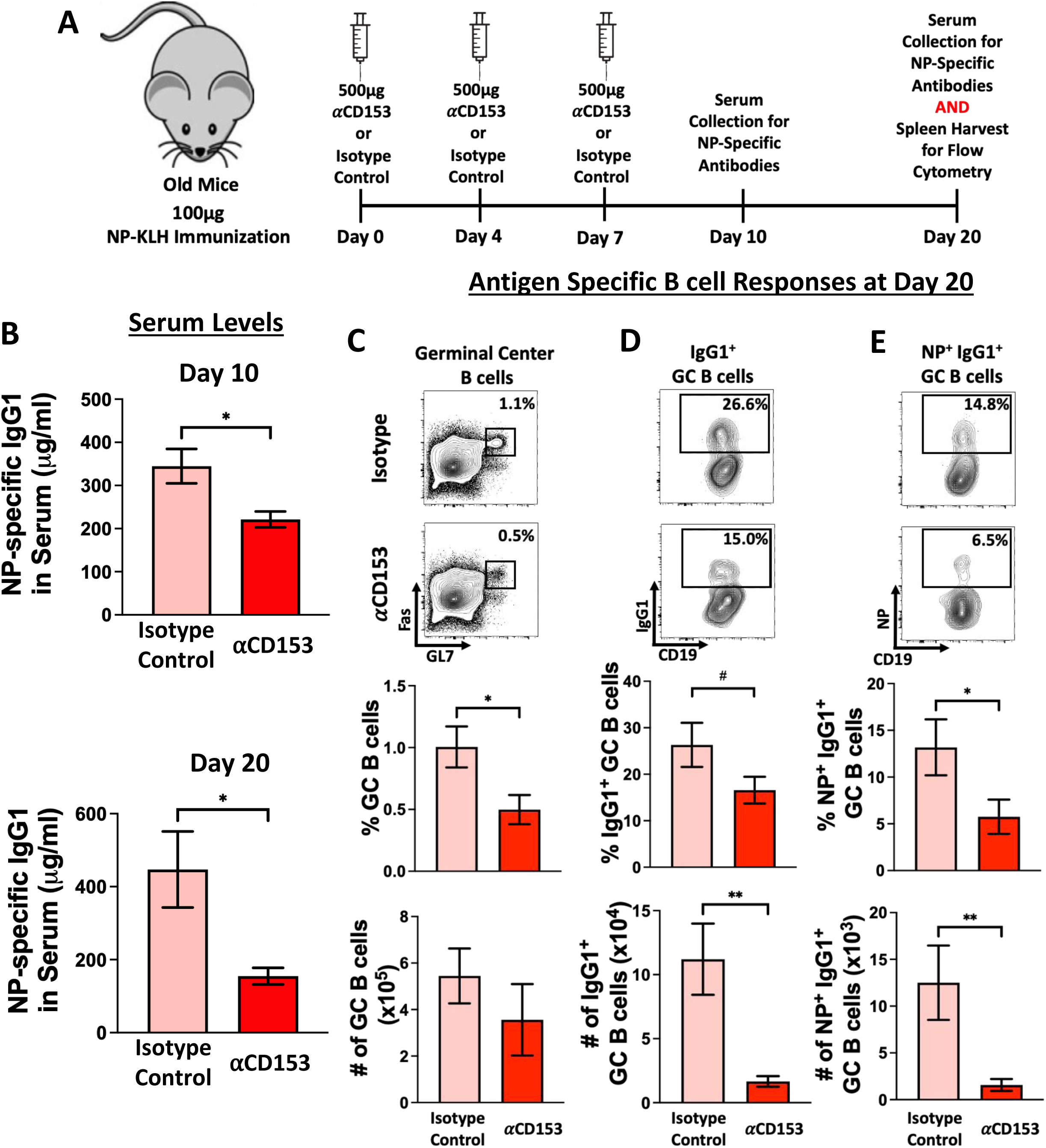
CD153 neutralization during immunization leads to decreased B cell responses. **A)** Aged WT (C57BL/6) mice were immunized with nitrophenol-keyhole limpet hemocyanin (NP-KLH) in alum and were give 500μg of either *α*CD153 neutralizing antibody or isotype control on day 0, 4, and 7. **B)** Serum levels of immunoglobulin G1 (IgG1) specific for NP on days 10 and 20 after immunization (n=5/group; mean±SEM). **C)** Plots and graphs display the frequency and number of germinal center B cells (gated on CD19^+^ B220^+^ Fas^+^ GL7^+^ cells), **D)** the frequency and number of IgG1+ GC B cells, and **E)** the frequency and number of NP-specific IgG1^+^ GC B cells (n=15-17/group with two replicates; mean±SEM). # *P* ≤ 0.08, \**P* ≤ 0.05, ** *P* ≤ 0.01, *** *P* ≤ 0.001, Student’s *t* test.

### CD153 neutralization decreases CD278 (ICOS) expression on antigen specific Tfh cells

As CD153 neutralization resulted in significantly decreased B cell responses, we next tested whether CD153 promoted Tfh cells after immunization. Aged wild-type mice were immunized with gp61-OVA and treated with a CD153 neutralizing antibody or an isotype control (Figure 7A). Neither overall frequency of total gp61-sp. cells nor their Tfh subset significantly changed in CD153 neutralized mice after 10 days (Figure 7B). Next, we determined whether CD153 blockade affected expression of co-stimulation molecules known to promote Tfh function, on antigen-specific Tfh cells. While expression of CD40L, OX40, CD30, 41BB, CD28, or GITR were not changed (Suppl. Figure 4A), the expression of CD278 (ICOS) was significantly decreased by CD153 neutralization (Figure 7C). ICOS is a costimulatory molecule present on Tfh cells that is vital for Tfh cell maintenance and T cell-dependent B cell responses such as immunoglobulin class-switching (*44, 45*). Thus, these data are consistent with a scenario in which CD153 promotes expression of ICOS, which is critical to promote sustained GC B cell responses.

**Figure 7:**
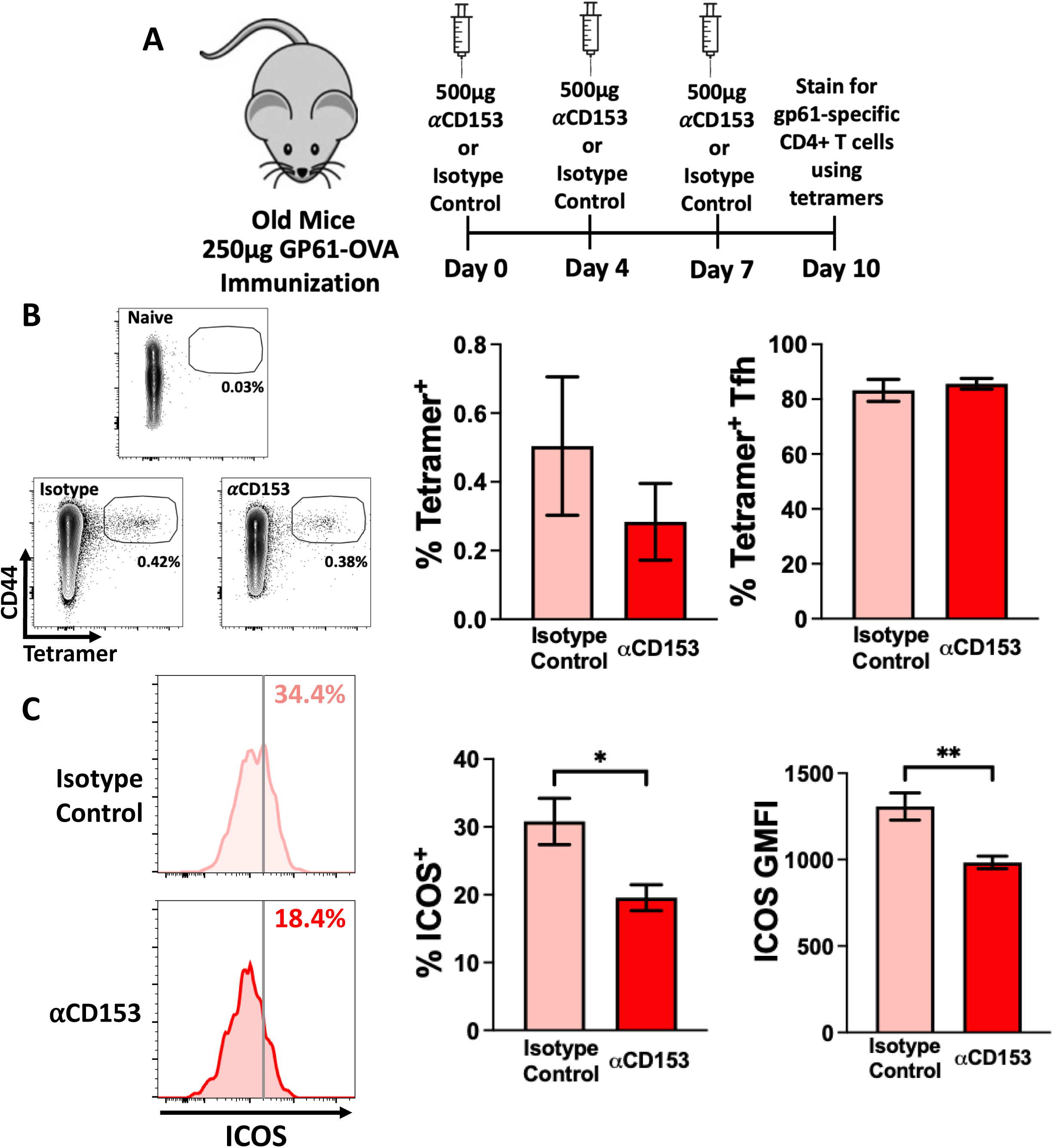
CD153 neutralization decreases (CD278) ICOS expression on antigen specific Tfh cells. **A)** Aged WT (C57BL/6) mice (n=5/group with two replicates) were immunized with 250μg of gp61 conjugated to OVA (gp61-OVA) in alum and were give 500μg of either *α*CD153 neutralizing antibody or isotype control on day 0, 4, and 7. Spleens were harvested on day 10 and analyzed by flow cytometry. **B)** Plots show IA^b^ -gp61 tetramer staining in naïve and immunized old mice that were treated with *α*CD153 neutralizing antibody or isotype control. Graphs show the frequency of gp61-specific total T cells and the frequency of Tfh cells (PD1^+^ CXCR5^+^) in the gp61-specifc T cell population (mean±SEM). **C)** Plots and graphs display the frequency of ICOS^+^ gp61-specific Tfh cells and the geometric mean fluorescence (GMFI) of ICOS in the gp61-specific Tfh cell population (mean±SEM). \**P* ≤ 0.05, ** *P* ≤ 0.01, *** *P* ≤ 0.001, Student’s *t* test.

## Discussion

Recent work, including our own, suggest age-dependent alternations in the Tfh compartment contribute to age-associated immune dysfunction and co-morbidities (*7, 13, 14, 25, 46*). In this study, we leveraged unbiased single-cell genomics profiling to transcriptionally characterize the memory CD4^+^ T cell compartment, including Tfh10 cells. Interestingly, our scRNAseq analyses highlighted CD153 as a marker of both the Tfh and Tfh10 cell populations. Increased CD153 expression on aged CD4^+^ T cells has been reported previously (*22-24, 47*), but has been attributed as a marker of “senescent” CD4^+^ T cells that accumulate with age. More than just a marker, a recent report showed that either CD153 or its receptor CD30 are required for accumulation of these “senescent” cells (*25*). In our companion study (*30*), genome-scale transcriptome comparisons show a one-to-one correspondence between these cells (*23*) and our Tfh10 population. Indeed, similar to Tfh10, the senescent cells express Tfh markers PD1, CXCR5, and lower levels of Bcl6 (*22, 48*). Interestingly, two recent studies have suggested that age-acquired CD4^+^ CD153^+^ memory T cells interact with age-associated B cells to sustain tertiary lymphoid structures in chronic kidney disease and produce spontaneous germinal center development in lupus (*22, 23*), also suggesting that these age-acquired CD153^+^ T cells have a Tfh cell phenotype. Our data are consistent with a functional role for these cells as we showed that neutralization of CD153 leads to decreased antigen specific germinal center B cells and antibody production after immunization. However, our data contrast a prior study showing that young CD153^-/-^ mice had a normal response to NP-OVA (*25*). A likely explanation for these differences is that aged Tfh cells rely more on CD153 than young Tfh cells, given their drastically increased expression of CD153.

CD153-expressing Tfh cells have both positive and negative impacts on aging-immune phenotypes. On the one hand, we show that they support the remaining capability to mount antibody responses to vaccination in old age. However, these cells can also be harmful in the context of autoimmune and chronic diseases associated with age. Regardless of their categorization as Tfh cells or senescent associated T cells, the emerging data clearly show that these cells are important for CD4^+^ T cell dependent immune response in aged mice, whether protective or pathologic. Therefore, targeting CD153 in autoimmunity as proposed (*23, 25*) may have unintended consequences (i.e., come at the cost of diminished responses to infection/vaccination in elderly individuals).

Mechanistically, our data show that CD153 contributes to the quality rather than the quantity of Tfh cells in response to immunization. Indeed, CD153 is important for normal ICOS expression on Tfh cells after vaccination. ICOS is critical for GC reactions (*49, 50*), including promotion of CD40L-dependent antibody class-switching (*51*) and a continual feed-forward loop in promoting T:B cell interactions in the germinal center (*52*). Thus, our data are consistent with a scenario where CD153 interactions promote or sustain late expression of ICOS which in turn, boosts B cell responses in the aged mice. However, further work is necessary to understand the underlying mechanism(s) of how CD153 expression impacts ICOS levels.

Importantly, our data also begins to shed light on the origins of elevated CD153 expression in aging. We show that CD153^+^ Tfh cells are retained after immunization in aged mice compared to younger mice. This suggests that the retention of antigen-specific CD153^+^ Tfh cells after immunological challenges may be one of the mechanisms for the accumulation of these cells with age. Given that us and others have reported increased levels of systemic IL-6 in aged mice (*7, 53*), and our data here show that IL-6 is both necessary and sufficient to drive expression of CD153, it is likely that CD4+ T cell responses generated in the presence of sustained high levels of IL-6 lead to the elevation and persistence of CD153 expression in aged mice. Further work is required to establish this relationship. Nonetheless, if true, this could be an important link between inflammaging and accumulation of CD153-expression Tfh cells.

Interestingly, while high levels of CD153 are expressed on all Tfh cells, its expression is even further elevated on IL-10-producing CD4^+^ T cells, including Tfh10 cells. While our previous data suggests that IL-10 inhibits B cell responses during vaccination in aged mice (*7*), here we report that CD153 promotes aged B cell responses. Thus, it is paradoxical that both IL-10 and CD153 expression are highly increased on the same cells, as they appear to play opposing roles in the aging environment. In our companion analysis of gene regulatory networks in the CD4+ memory T cell compartment (*30*), we identify several transcription factors (e.g. Irf4, Maf, Nfia, Nfat1/3, Vdr) that collectively promote Tfh10 gene signatures of contrasting effector (e.g., CD153) and regulatory (e.g., IL-10) genes. Here, we establish c-Maf as an important regulator of Tfh10 cells and CD153 expression. Other groups have reported a role for c-Maf in control of IL-10 expression in CD4^+^ T cells (*42, 54, 55*). Thus, this collective evidence points to c-Maf as a critical regulator of both molecules and likely other inhibitory and stimulatory genes expressed by Tfh cells. Future work will determine how c-Maf collaborates with other age-dependent transcription factors to mediate its diverse functions in aging T cells.

Overall, our data suggests a role for the accumulation and function of CD153^+^ CD4^+^ T cells during aging. Our data connect IL-6 and inflammaging to accumulation of not only Tfh10 cells in our prior work but also CD153^+^ Tfh cells in this work. Further, we found that expression of CD153 is vital in the context of vaccination, as blocking its interaction diminishes B cell responses after immunization in aged mice. Thus, additional work is necessary to understand the potential protective (via CD153) or detrimental (via IL-10) roles for these cells on vaccine responsiveness. Understanding the mechanisms underlying protective versus detrimental roles of these cells could be exploited to enhance vaccine responses while limiting auto- or allo-immune antibody responses.

## ACKNOWLEDGEMENTS

The authors declare they have no competing interests and they thank the members of the Hildeman, Chougnet, and Miraldi labs for their helpful discussion. All data needed to evaluate the conclusions in the paper are present in the paper and/or the Supplementary Materials. This research was also made possible, in part, using the Cincinnati Children’s Single Cell Genomics Core, the DNA Sequencing and Genotyping Core, and Biomedical Informatics Core. We specifically acknowledge the assistance of Kelly Rangel and Shawn Smith from the Single Cell Genomics Core. This work was supported by Public Health Service grants AG033057 (D.A.H. and C.A.C.), AG033057 (D.A.H. and C.A.C.), AG053498 (D.A.H. and C.A.C.), U01AI150748 (E.R.M.), R01AI153442 (E.R.M.), and R21AI156185 (E.R.M.). All flow cytometric data were acquired using equipment maintained by the Research Flow Cytometry Core in the Division of Rheumatology at Cincinnati Children’s Hospital Medical Center which is supported in part by NIH AR070549 and by the Hematology Center of Excellence U54DK126108.

## References

1. J. M. Lord, The effect of aging of the immune system on vaccination responses. Human vaccines & immunotherapeutics 9, 1364–1367 (2013).

2. J. J. Goronzy, F. Fang, M. M. Cavanagh, Q. Qi, C. M. Weyand, Naive T cell maintenance and function in human aging. The Journal of Immunology 194, 4073–4080 (2015).

3. M. A. Moro-García, R. Alonso-Arias, C. López-Larrea, When aging reaches CD4+ T-cells: phenotypic and functional changes. Frontiers in immunology 4, 107 (2013).

4. J. J. Goronzy, C. M. Weyand, Successful and maladaptive T cell aging. Immunity 46, 364–378 (2017).

5. M. Mittelbrunn, G. Kroemer, Hallmarks of T cell aging. Nature immunology 22, 687–698 (2021).

6. S. Crotty, T follicular helper cell biology: a decade of discovery and diseases. Immunity 50, 1132–1148 (2019).

7. M. Almanan, J. Raynor, I. Ogunsulire, A. Malyshkina, S. Mukherjee, S. A. Hummel, J. T. Ingram, A. Saini, M. M. Xie, T. Alenghat, IL-10–producing Tfh cells accumulate with age and link inflammation with age-related immune suppression. Science advances 6, eabb0806 (2020).

8. M. E. Weksler, P. Szabo, The effect of age on the B-cell repertoire. Journal of clinical immunology 20, 240–249 (2000).

9. B. Zheng, S. Han, Y. Takahashi, G. Kelsoe, Immunosenescence and germinal center reaction. Immunological reviews 160, 63–77 (1997).

10. H. Song, P. W. Price, J. Cerny, Age-related changes in antibody repertoire: contribution from T cells. Immunological reviews 160, 55–62 (1997).

11. M. Kosco, G. Burton, Z. Kapasi, A. Szakal, J. Tew, Antibody-forming cell induction during an early phase of germinal centre development and its delay with ageing. Immunology 68, 312 (1989).

12. A. K. Szakal, J. K. Taylor, J. P. Smith, M. H. Kosco, G. F. Burton, J. J. Tew, Kinetics of germinal center development in lymph nodes of young and aging immune mice. The Anatomical Record 227, 475–485 (1990).

13. J. S. Lefebvre, E. C. Lorenzo, A. R. Masters, J. W. Hopkins, S. M. Eaton, S. T. Smiley, L. Haynes, Vaccine efficacy and T helper cell differentiation change with aging. Oncotarget 7, 33581 (2016).

14. P. T. Sage, C. L. Tan, G. J. Freeman, M. Haigis, A. H. Sharpe, Defective TFH cell function and increased TFR cells contribute to defective antibody production in aging. Cell reports 12, 163–171 (2015).

15. M. Yu, G. Li, W.-W. Lee, M. Yuan, D. Cui, C. M. Weyand, J. J. Goronzy, Signal inhibition by the dual-specific phosphatase 4 impairs T cell-dependent B-cell responses with age. Proceedings of the National Academy of Sciences 109, E879–E888 (2012).

16. S. Han, K. Hathcock, B. Zheng, T. B. Kepler, R. Hodes, G. Kelsoe, Cellular interaction in germinal centers. Roles of CD40 ligand and B7-2 in established germinal centers. The Journal of Immunology 155, 556–567 (1995).

17. E. L. Stone, M. Pepper, C. D. Katayama, Y. M. Kerdiles, C.-Y. Lai, E. Emslie, Y. C. Lin, E. Yang, A. W. Goldrath, M. O. Li, ICOS coreceptor signaling inactivates the transcription factor FOXO1 to promote Tfh cell differentiation. Immunity 42, 239–251 (2015).

18. D. Liu, H. Xu, C. Shih, Z. Wan, X. Ma, W. Ma, D. Luo, H. Qi, T–B-cell entanglement and ICOSL-driven feed-forward regulation of germinal centre reaction. Nature 517, 214–218 (2015).

19. E. K. Deenick, C. S. Ma, R. Brink, S. G. Tangye, Regulation of T follicular helper cell formation and function by antigen presenting cells. Current opinion in immunology 23, 111–118 (2011).

20. E. Perkey, R. A. Miller, G. G. Garcia, Ex vivo enzymatic treatment of aged CD4 T cells restores cognate T cell helper function and enhances antibody production in mice. The Journal of Immunology 189, 5582–5589 (2012).

21. S. M. Eaton, E. M. Burns, K. Kusser, T. D. Randall, L. Haynes, Age-related defects in CD4 T cell cognate helper function lead to reductions in humoral responses. The Journal of experimental medicine 200, 1613–1622 (2004).

22. Y. Fukushima, N. Minato, M. Hattori, The impact of senescence-associated T cells on immunosenescence and age-related disorders. Inflammation and Regeneration 38, 1–6 (2018).

23. Y. Sato, A. Oguchi, Y. Fukushima, K. Masuda, N. Toriu, K. Taniguchi, T. Yoshikawa, X. Cui, M. Kondo, T. Hosoi, S. Komidori, Y. Shimizu, H. Fujita, L. Jiang, Y. Kong, T. Yamanashi, J. Seita, T. Yamamoto, S. Toyokuni, Y. Hamazaki, M. Hattori, Y. Yoshikai, P. Boor, J. Floege, H. Kawamoto, Y. Murakawa, N. Minato, M. Yanagita, CD153/CD30 signaling promotes age-dependent tertiary lymphoid tissue expansion and kidney injury. J Clin Invest 132, (2022).

24. S. Tahir, Y. Fukushima, K. Sakamoto, K. Sato, H. Fujita, J. Inoue, T. Uede, Y. Hamazaki, M. Hattori, N. Minato, A CD153+ CD4+ T follicular cell population with cell-senescence features plays a crucial role in lupus pathogenesis via osteopontin production. The Journal of Immunology 194, 5725–5735 (2015).

25. Y. Fukushima, K. Sakamoto, M. Matsuda, Y. Yoshikai, H. Yagita, D. Kitamura, M. Chihara, N. Minato, M. Hattori, cis interaction of CD153 with TCR/CD3 is crucial for the pathogenic activation of senescence-associated T cells. Cell Reports 40, 111373 (2022).

26. K. D. Shanebeck, C. R. Maliszewski, M. K. Kennedy, K. S. Picha, C. A. Smith, R. G. Goodwin, K. H. Grabstein, Regulation of murine B cell growth and differentiation by CD30 ligand. European journal of immunology 25, 2147–2153 (1995).

27. F. M. Gaspal, M.-Y. Kim, F. M. McConnell, C. Raykundalia, V. Bekiaris, P. J. Lane, Mice deficient in OX40 and CD30 signals lack memory antibody responses because of deficient CD4 T cell memory. The Journal of Immunology 174, 3891–3896 (2005).

28. M. D. Jumper, Y. Nishioka, L. S. Davis, P. E. Lipsky, K. Meek, Regulation of human B cell function by recombinant CD40 ligand and other TNF-related ligands. The Journal of Immunology 155, 2369–2378 (1995).

29. A. Cerutti, A. Schaffer, S. Shah, H. Zan, H.-C. Liou, R. G. Goodwin, P. Casali, CD30 is a CD40inducible molecule that negatively regulates CD40-mediated immunoglobulin class switching in non-antigen-selected human B cells. Immunity 9, 247–256 (1998).

30. J. A. Wayman, A. Thomas, A. Bejjani, A. Katko, M. Almanan, A. Godarova, S. Korinfskaya, T. A. Cazares, M. Yukawa, L. C. Kottyan, A. Barski, C. A. Chougnet, D. A. Hildeman, E. R. Miraldi, An atlas of gene regulatory networks for T memory cells in youth and old age. bioRxiv, 2023.2003.2007.531590 (2023).

31. Y. Hao, S. Hao, E. Andersen-Nissen, W. M. Mauck, 3rd, S. Zheng, A. Butler, M. J. Lee, A. J. Wilk, C. Darby, M. Zager, P. Hoffman, M. Stoeckius, E. Papalexi, E. P. Mimitou, J. Jain, A. Srivastava, T. Stuart, L. M. Fleming, B. Yeung, A. J. Rogers, J. M. McElrath, C. A. Blish, R. Gottardo, P. Smibert, R. Satija, Integrated analysis of multimodal single-cell data. Cell 184, 3573-3587.e3529 (2021).

32. C. S. McGinnis, L. M. Murrow, Z. J. Gartner, DoubletFinder: doublet detection in single-cell RNA sequencing data using artificial nearest neighbors. Cell systems 8, 329-337. e324 (2019).

33. T. Stuart, A. Butler, P. Hoffman, C. Hafemeister, E. Papalexi, W. M. Mauck III, Y. Hao, M. Stoeckius, P. Smibert, R. Satija, Comprehensive integration of single-cell data. Cell 177, 1888-1902. e1821 (2019).

34. F. Andris, S. Denanglaire, M. Anciaux, M. Hercor, H. Hussein, O. Leo, The transcription factor c-Maf promotes the differentiation of follicular helper T cells. Frontiers in immunology 8, 480 (2017).

35. Y. Hiramatsu, A. Suto, D. Kashiwakuma, H. Kanari, S. i. Kagami, K. Ikeda, K. Hirose, N. Watanabe, M. J. Grusby, I. Iwamoto, c-Maf activates the promoter and enhancer of the IL-21 gene, and TGF-β inhibits c-Maf-induced IL-21 production in CD4+ T cells. Journal of leukocyte biology 87, 703–712 (2010).

36. A. Sahoo, A. Alekseev, K. Tanaka, L. Obertas, B. Lerman, C. Haymaker, K. Clise-Dwyer, J. S. McMurray, R. Nurieva, Batf is important for IL-4 expression in T follicular helper cells. Nature communications 6, 1–10 (2015).

37. H. Zhang, A. Madi, N. Yosef, N. Chihara, A. Awasthi, C. Pot, C. Lambden, A. Srivastava, P. R. Burkett, J. Nyman, E. Christian, Y. Etminan, A. Lee, H. Stroh, J. Xia, K. Karwacz, P. I. Thakore, N. Acharya, A. Schnell, C. Wang, L. Apetoh, O. Rozenblatt-Rosen, A. C. Anderson, A. Regev, V. K. Kuchroo, An IL-27-Driven Transcriptional Network Identifies Regulators of IL-10 Expression across T Helper Cell Subsets. Cell Rep 33, 108433 (2020).

38. M. Ciofani, A. Madar, C. Galan, M. Sellars, K. Mace, F. Pauli, A. Agarwal, W. Huang, C. N. Parkhurst, M. Muratet, K. M. Newberry, S. Meadows, A. Greenfield, Y. Yang, P. Jain, F. K. Kirigin, C. Birchmeier, E. F. Wagner, K. M. Murphy, R. M. Myers, R. Bonneau, D. R. Littman, A validated regulatory network for Th17 cell specification. Cell 151, 289–303 (2012).

39. J. Raynor, R. Karns, M. Almanan, K.-P. Li, S. Divanovic, C. A. Chougnet, D. A. Hildeman, IL-6 and ICOS antagonize bim and promote regulatory T cell accrual with age. The Journal of Immunology 195, 944–952 (2015).

40. R. Daynes, B. Araneo, W. Ershler, C. Maloney, G.-Z. Li, S.-Y. Ryu, Altered regulation of IL-6 production with normal aging. Possible linkage to the age-associated decline in dehydroepiandrosterone and its sulfated derivative. The Journal of Immunology 150, 5219–5230 (1993).

41. M. Jergović, H. L. Thompson, C. M. Bradshaw, S. A. Sonar, A. Ashgar, N. Mohty, B. Joseph, M. J. Fain, K. Cleveland, R. G. Schnellman, J. Nikolich-Žugich, IL-6 can singlehandedly drive many features of frailty in mice. Geroscience 43, 539–549 (2021).

42. Y. Yang, J. Ochando, A. Yopp, J. S. Bromberg, Y. Ding, IL-6 plays a unique role in initiating c-Maf expression during early stage of CD4 T cell activation. The Journal of Immunology 174, 2720–2729 (2005).

43. M. Liu, Z. Tong, C. Ding, F. Luo, S. Wu, C. Wu, S. Albeituni, L. He, X. Hu, D. Tieri, Transcription factor c-Maf is a checkpoint that programs macrophages in lung cancer. The Journal of clinical investigation 130, 2081–2096 (2020).

44. A. Tafuri, A. Shahinian, F. Bladt, S. K. Yoshinaga, M. Jordana, A. Wakeham, L.-M. Boucher, D. Bouchard, V. S. Chan, G. Duncan, ICOS is essential for effective T-helper-cell responses. Nature 409, 105–109 (2001).

45. H. Akiba, K. Takeda, Y. Kojima, Y. Usui, N. Harada, T. Yamazaki, J. Ma, K. Tezuka, H. Yagita, K. Okumura, The role of ICOS in the CXCR5+ follicular B helper T cell maintenance in vivo. The Journal of Immunology 175, 2340–2348 (2005).

46. D. L. Hill, C. E. Whyte, S. Innocentin, J. L. Lee, J. Dooley, J. Wang, E. A. James, J. C. Lee, W. W. Kwok, M. S. Zand, A. Liston, E. J. Carr, M. A. Linterman, Impaired HA-specific T follicular helper cell and antibody responses to influenza vaccination are linked to inflammation in humans. Elife 10, (2021).

47. S. Yoshida, H. Nakagami, H. Hayashi, Y. Ikeda, J. Sun, A. Tenma, H. Tomioka, T. Kawano, M. Shimamura, R. Morishita, The CD153 vaccine is a senotherapeutic option for preventing the accumulation of senescent T cells in mice. Nature communications 11, 1–10 (2020).

48. K. Sakamoto, Y. Fukushima, K. Ito, M. Matsuda, S. Nagata, N. Minato, M. Hattori, Osteopontin in spontaneous germinal centers inhibits apoptotic cell engulfment and promotes antinuclear antibody production in lupus-prone mice. The Journal of Immunology 197, 2177–2186 (2016).

49. C. Dong, U.-A. Temann, R. A. Flavell, Cutting edge: critical role of inducible costimulator in germinal center reactions. The Journal of Immunology 166, 3659–3662 (2001).

50. L. Bossaller, J. Burger, R. Draeger, B. Grimbacher, R. Knoth, A. Plebani, A. Durandy, U. Baumann, M. Schlesier, A. A. Welcher, ICOS deficiency is associated with a severe reduction of CXCR5+ CD4 germinal center Th cells. The Journal of Immunology 177, 4927–4932 (2006).

51. A. J. McAdam, R. J. Greenwald, M. A. Levin, T. Chernova, N. Malenkovich, V. Ling, G. J. Freeman, A. H. Sharpe, ICOS is critical for CD40-mediated antibody class switching. Nature 409, 102–105 (2001).

52. V. Panneton, J. Chang, M. Witalis, J. Li, W. K. Suh, Inducible T-cell co-stimulator: Signaling mechanisms in T follicular helper cells and beyond. Immunological Reviews 291, 91–103 (2019).

53. C. Franceschi, M. Bonafè, S. Valensin, F. Olivieri, M. De Luca, E. Ottaviani, G. De Benedictis, Inflamm-aging: an evolutionary perspective on immunosenescence. Annals of the new York Academy of Sciences 908, 244–254 (2000).

54. J. Xu, Y. Yang, G. Qiu, G. Lal, Z. Wu, D. E. Levy, J. C. Ochando, J. S. Bromberg, Y. Ding, c-Maf regulates IL-10 expression during Th17 polarization. The Journal of Immunology 182, 6226–6236 (2009).

55. C. Neumann, F. Heinrich, K. Neumann, V. Junghans, M.-F. Mashreghi, J. Ahlers, M. Janke, C. Rudolph, N. Mockel-Tenbrinck, A. A. Kühl, Role of Blimp-1 in programing Th effector cells into IL-10 producers. Journal of Experimental Medicine 211, 1807–1819 (2014).

56. C. E. Grant, T. L. Bailey, W. S. Noble, FIMO: scanning for occurrences of a given motif. Bioinformatics 27, 1017–1018 (2011).

