## Supplemental figures for "Elevated CD153 Expression on Aged T Follicular Helper Cells is Vital for B cell Responses"

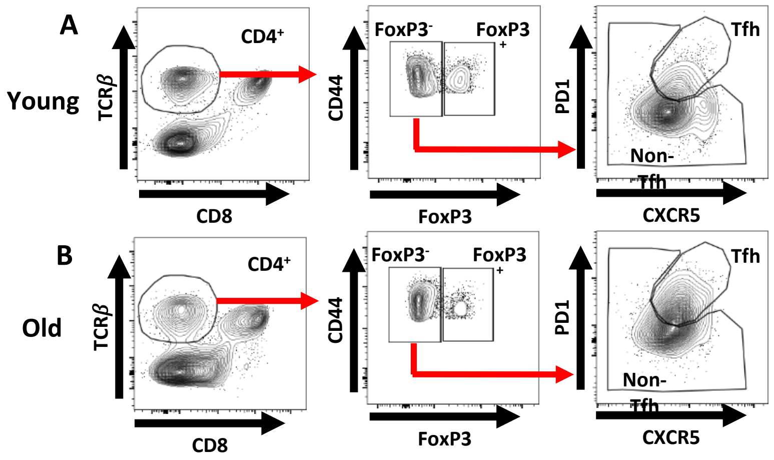


**Supplementary Figure 1: Flow cytometry gating strategy to identify Tregs, non-Tfh cells, and Tfh cells.** Representative flow cytometry gating of A) young and B) old mouse spleens to identify Tregs ($CD4^+$ $FoxP3^+$), non-Tfh cells ($CD4^+$ $FoxP3^-$ $PD1^-$ $CXCR5^-$), and Tfh cells ($CD4^+$ $FoxP3^-$ $PD1^+$ $CXCR5^+$).

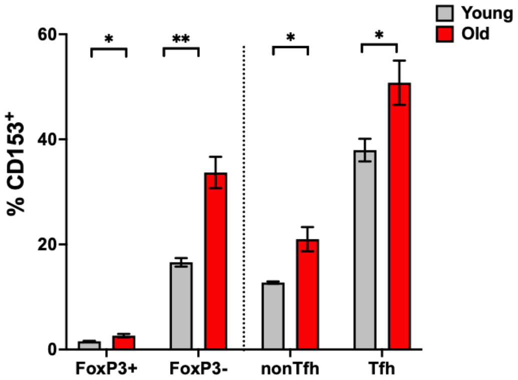


**Supplementary Figure 2: CD153 expression in CD4$^+$ memory T cell compartments.** Splenocytes of young and aged C57BL/6 (n=4/group) were stimulated with P + I, stained with antibodies against CD44, TCRβ, CD8, FoxP3, IL-10, and CD153, and analyzed by flow cytometry. Graph shows the frequency of CD153$^+$ cells in the CD4$^+$ T cell memory compartment (CD44$^+$TCRβ$^+$ CD8$^-$) (means±SEM). *$P \leq 0.05$, ** $P \leq 0.01$, *** $P \leq 0.001$, Student's *t* test.

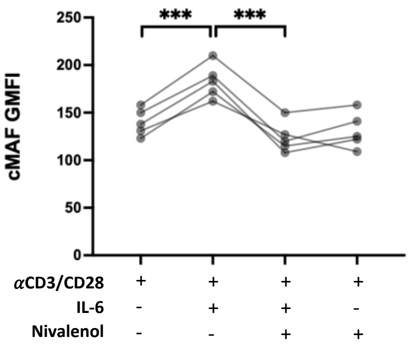


**Supplementary Figure 3: c-Maf expression is modulated by IL-6 and Nivalenol.** CD4+ T cells were isolated from young WT mice and stimulated *in vitro* with αCD3/αCD28 in the presence or absence of IL-6 and nivalenol. Graph shows the gMFI of c-Maf in CD4+ T cells after 24 hours. Each line represents one mouse across treatment groups. *$P \leq 0.05$, ** $P \leq 0.01$, *** $P \leq 0.001$, Student's *t* test.

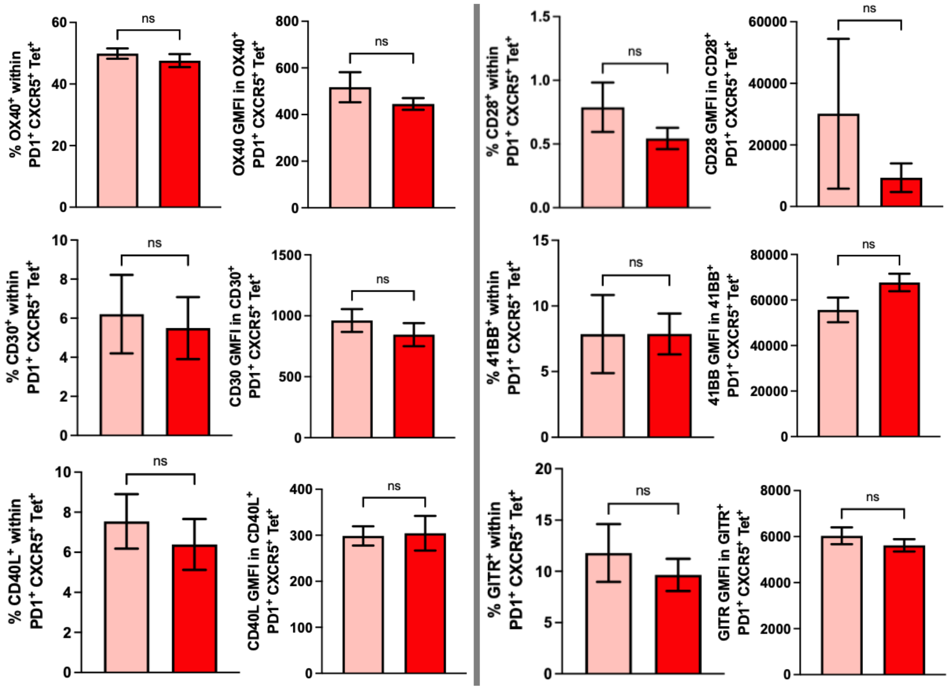


**Supplementary Figure 4: CD153 neutralization did not affect antigen-specific Tfh cell expression of co-stimulatory molecules other than ICOS.** Aged WT (C57BL/6) mice (n=5 per group) were immunized with 250µg of gp61 conjugated to OVA (gp61-OVA) in alum and were give 500µg of either $\alpha$CD153 neutralizing antibody or isotype control on day 0, 4, and 7. Spleens were harvested on day 10 and analyzed by flow cytometry for costimulatory molecule expression. Plots and graphs display the frequency of gp61-specific Tfh cells positive for costimulatory markers and gMFI of those markers within the positive population (means±SEM **P* ≤ 0.05, ** *P* ≤ 0.01, *** *P* ≤ 0.001, Student's *t* test.
